## Supplementary material for "Tuning social interactions’ strength drives collective response to light intensity in schooling fish": https://www.dropbox.com/s/uv728ekkknyev0b/SM_Article_TingtingXue_et_al.pdf?dl=0

### Supplementary Information for
Tuning social interactions strength drives collective response to light intensity in schooling fish

Xue *et al.*

#### This PDF file includes:

Details of numerical simulations for preparing the figures of the manuscript and additional supporting results are presented here.
Supplementary Figures 1 to 13.
Supplementary Tables 1 to 8.
Supplementary Movies 1 to 8.

#### Other Supplementary Materials for this manuscript include the following:

Supplementary Movies 1 to 8
Supplementary Data 1

### Supplementary Figures

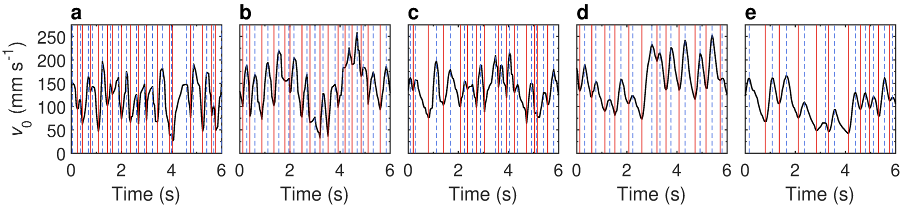


**Supplementary Fig. 1 Burst-and-coast motion of fish swimming alone in the tank.** Time series of the instantaneous speed of one fish under different light intensities: **a** 0.5, **b** 1, **c** 1.5, **d** 5 and **e** 50 lx. Colored vertical lines represent local minima (red) and maxima (blue) of the speed. Time intervals going from a red line to the next blue line correspond to the bursting acceleration phase, intervals going from a blue line to the next red line correspond to the decelerating gliding phase.

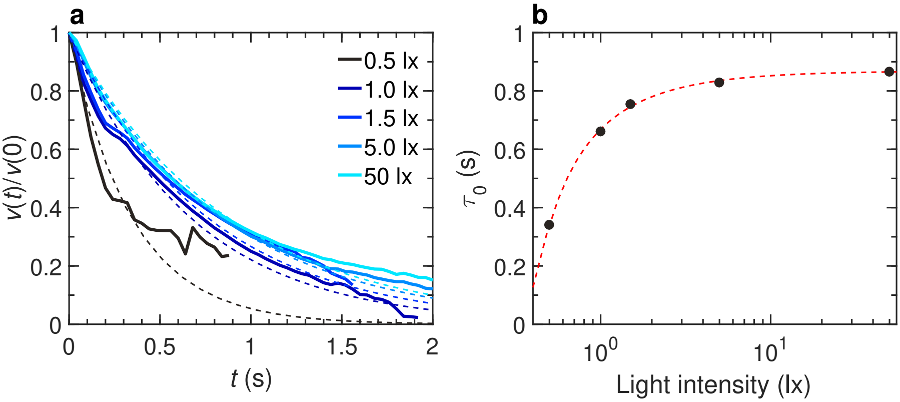


**Supplementary Fig. 2 Normalized average decay of fish speed right after a kick when the fish swims alone ($N = 1$). a** Exponential deceleration during the gliding phase averaged along all kicks and normalized with the value of the speed at the kicking instant, for different light intensities 0, 0.5, 1, 5, and 50 lx (from dark to light blue). Wide solid lines are experimental measures, dashed lines are exponential approximations of the form $\exp(-t/\tau_0)$, where $\tau_0$ is the relaxation time: $\tau_0 \approx 0.34$ (0.5 lx), 0.66 (1 lx), 0.76 (1.5 lx), 0.83 (5 lx), 0.87 (50 lx). **b** Mean relaxation time $\tau_0$ as a function of the light intensity (black circles). The red dashed line shows the trend of the average value with the light intensity.

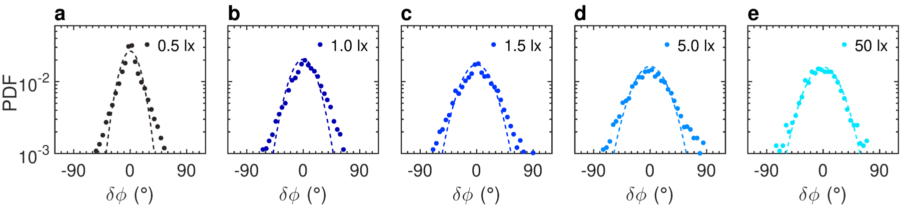


**Supplementary Fig. 3 Effects of light intensity on the spontaneous heading change of a fish swimming alone ($N = 1$).** Probability density function (PDF) of the angle variation $\delta\phi$ when the fish is far from the wall ($r_\mathrm{w} > 60$ mm) in five different light intensities: 0.5, 1, 1.5, 5 and 50 lx (from dark to light blue). Colored dots: measures from the experiments. Solid lines: approximation with Gaussian distributions, with $\gamma_R$=0.26, 0.34, 0.40, 0.42, and 0.43 respectively.

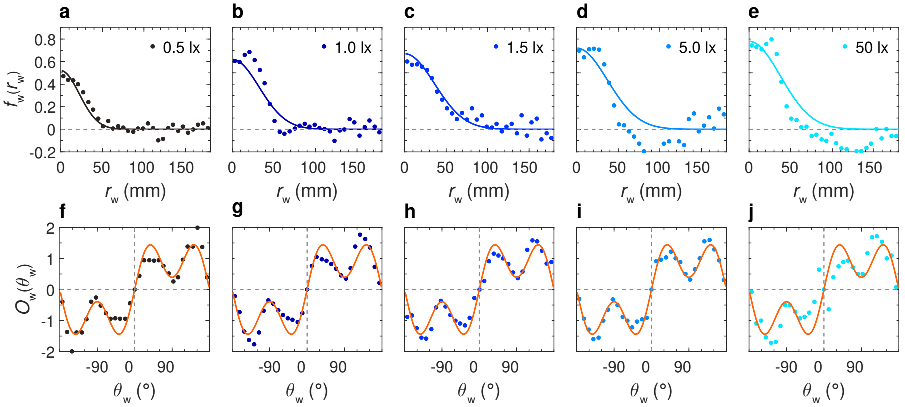


**Supplementary Fig. 4 Effect of light intensity on the fish interaction with the tank wall when the fish swims alone ($N = 1$).** Function of repulsion $f_\mathrm{w}(r_\mathrm{w})O_\mathrm{w}(\theta_\mathrm{w})$ as extracted from the experiments by means of the reconstruction procedure (dots), and analytical approximations used in the numerical simulations (solid lines), for different light intensities: 0.5, 1, 1.5, 5, and 50 lx (from dark to light blue). **a-e** Intensity of the interaction $f_\mathrm{w}(r_\mathrm{w})$ as a function of the fish distance to the wall $r_\mathrm{w}$. **f-j** Intensity of the interaction $O_\mathrm{w}(\theta_\mathrm{w})$ as a function of the relative orientation of the fish to the wall $\theta_\mathrm{w}$. Orange lines correspond to the analytical approximation of a single discrete function combining all light conditions: $O_\mathrm{w}(\theta_\mathrm{w}) = 1.9612\sin(\theta_\mathrm{w})[1 + 0.8\cos(2\theta_\mathrm{w})]$.

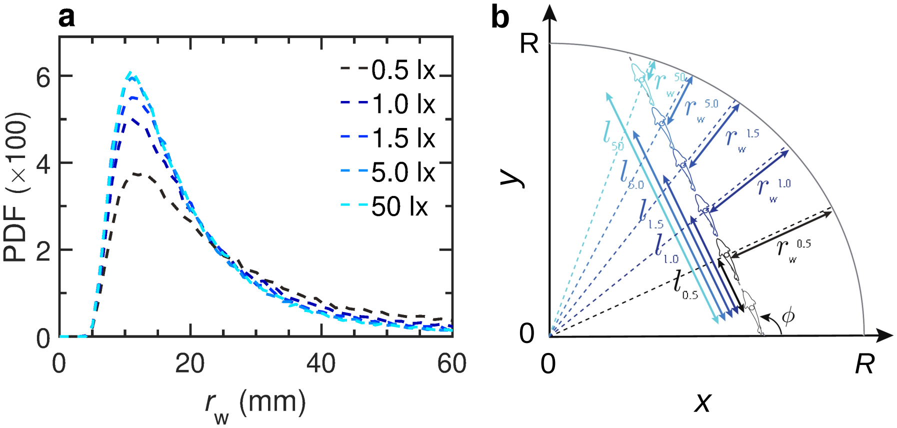


**Supplementary Fig. 5 Effect of the kick length $l$ on the spatial distribution of a fish swimming alone ($N = 1$).** **a** Probability density function (PDF) of the distance of the fish to the wall $r_{\mathrm{w}}$ as a function of light intensity when only the kick length $l$ is changed in the model. **b** Schematic diagram of the motion of a single fish between two kicks. The distance travelled by the fish between two kicks is greater at 50 lx than at 0.5 lx; as a consequence, the fish moves closer to the wall when light intensity is higher.

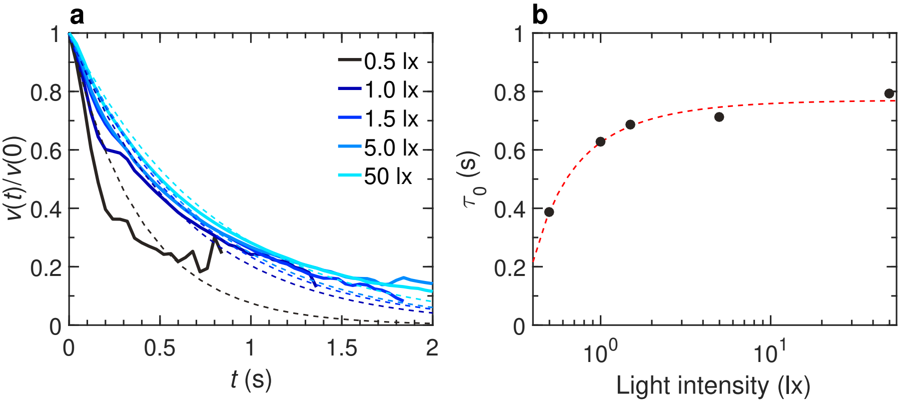


**Supplementary Fig. 6 Normalized average decay of fish speed right after a kick when fish swim in pairs ($N = 2$).** **a** Exponential deceleration during the gliding phase averaged along all kicks and normalized with the value of the speed at the kicking instant, for different light intensities 0, 0.5, 1, 5, and 50 lx (from dark to light blue). Wide solid lines are experimental measures, dashed lines are exponential approximations of the form $\exp(-t/\tau_0)$, where $\tau_0$ is the relaxation time: $\tau_0 \approx 0.39$ (0.5 lx), 0.63 (1 lx), 0.69 (1.5 lx), 0.71 (5 lx), 0.79 (50 lx). **b** Mean relaxation time $\tau_0$ as a function of the light intensity (black circles). The red dashed line shows the trend of the average value with the light intensity.

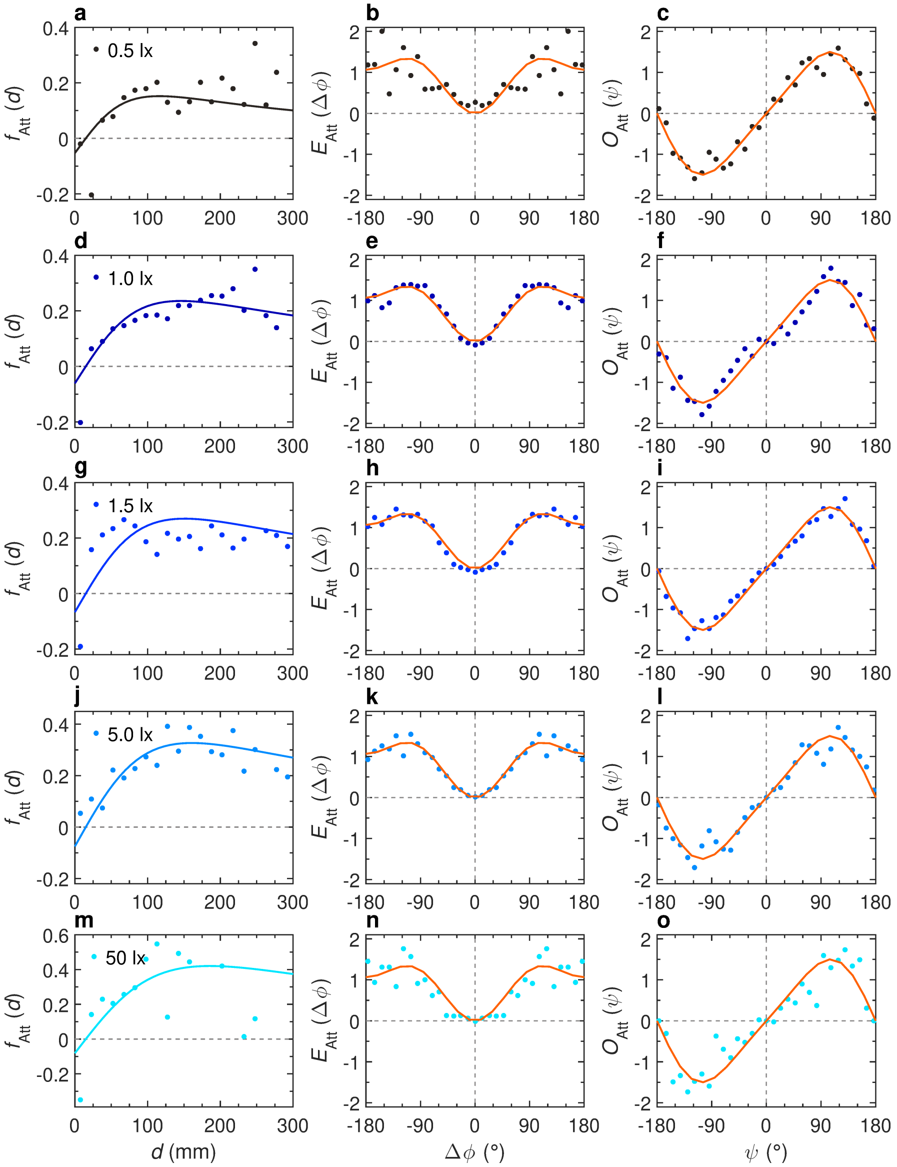


**Supplementary Fig. 7 Effects of light intensity on the attraction interaction between two fish ($N = 2$).** Components of the attraction interaction function $f_{\mathrm{Att}}(d)$, $O_{\mathrm{Att}}(\psi)$, and $E_{\mathrm{Att}}(\Delta\phi)$ as functions of the distance between fish $d$, the viewing angle $\psi$, and the relative heading $\Delta\phi$, for different light intensities: **a-c** 0.5 lx, **d-f** 1 lx, **g-i** 1.5 lx, **j-l** 5 lx, and **m-o** 50 lx (from dark to light blue). Color dots correspond to the discrete values resulting from the reconstruction procedure, extracted from the experimental data of the corresponding intensity of light. Solid lines correspond to the analytical approximation of the discrete function. Orange lines correspond to the analytical approximation of a single discrete function combining all light conditions.

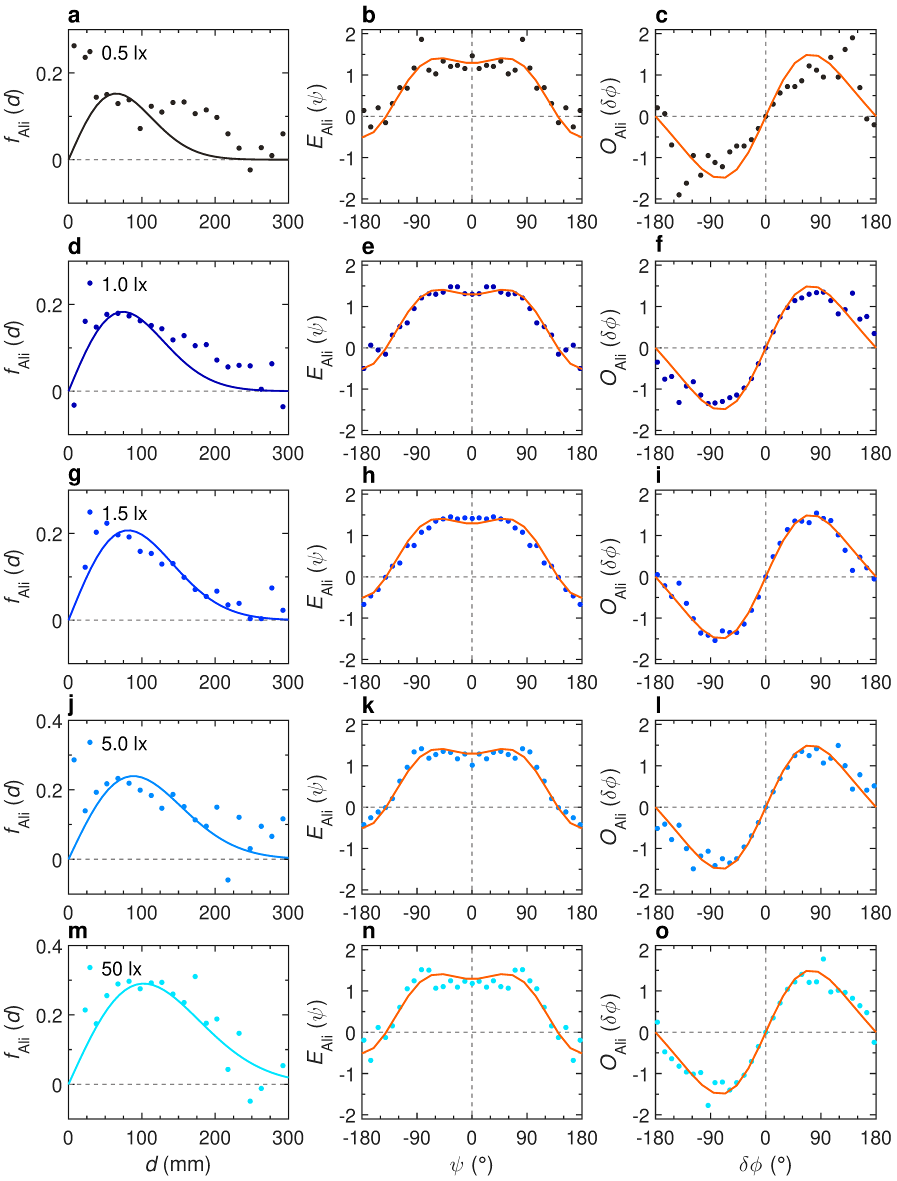


**Supplementary Fig. 8 Effects of light intensity on the alignment interaction between two fish ($N = 2$)** Components of the attraction interaction function $f_{\mathrm{Ali}}(d)$, $E_{\mathrm{Ali}}(\psi)$, and $O_{\mathrm{Ali}}(\Delta\phi)$ as functions of the distance between fish $d$, the viewing angle $\psi$, and the relative heading $\Delta\phi$, for different light intensities: **a-c** 0.5 lx, **d-f** 1 lx, **g-i** 1.5 lx, **j-l** 5 lx, and **m-o** 50 lx (from dark to light blue). Color dots correspond to the discrete values resulting from the reconstruction procedure, extracted from the experimental data of the corresponding intensity of light. Solid lines correspond to the analytical approximation of the discrete function. Orange lines correspond to the analytical approximation of a single discrete function combining all light conditions.

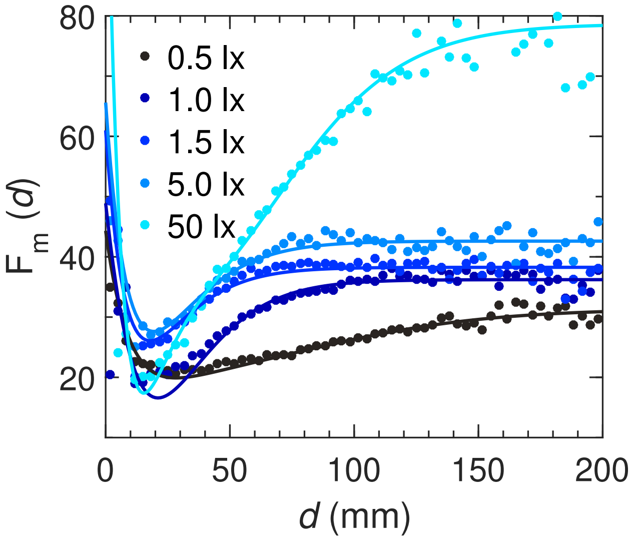


**Supplementary Fig. 9 Effect of light intensity on the modulation of the kick length with the distance between fish**. Modulation function $F_\mathrm{m}(d)$ of the mean value used in the distribution from which kick lengths are sampled, as a function of the distance between fish $d$, and for different light intensities: 0.5, 1, 1.5, 5, and 50 lx (from dark to light blue). Dots correspond to the discrete functions resulting from the reconstruction procedure and extracted from the experimental data. Solid lines correspond to the smooth analytical approximations of these discrete functions.

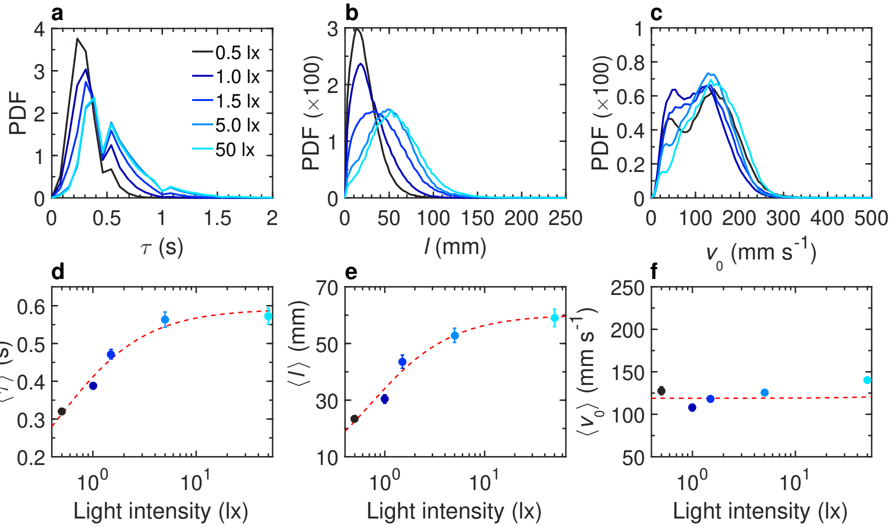


**Supplementary Fig. 10 Effects of light intensity on burst-and-coast swimming in groups of 5 fish. a-c** Probability density function (PDF) of kick duration $\tau$, kick length $l$, and peak speed $v_0$ respectively, at different light intensities: 0.5, 1, 1.5, 5 and 50 lx (from dark to light blue). **d-f** Average value of kick duration $\langle\tau\rangle$, kick length $\langle l\rangle$, and peak speed $\langle v_0\rangle$ respectively, at different light intensities. Solid circles are the average values on all experiments; error bars represent the standard deviation. Red dashed lines show the trend of the average value with the light intensity.

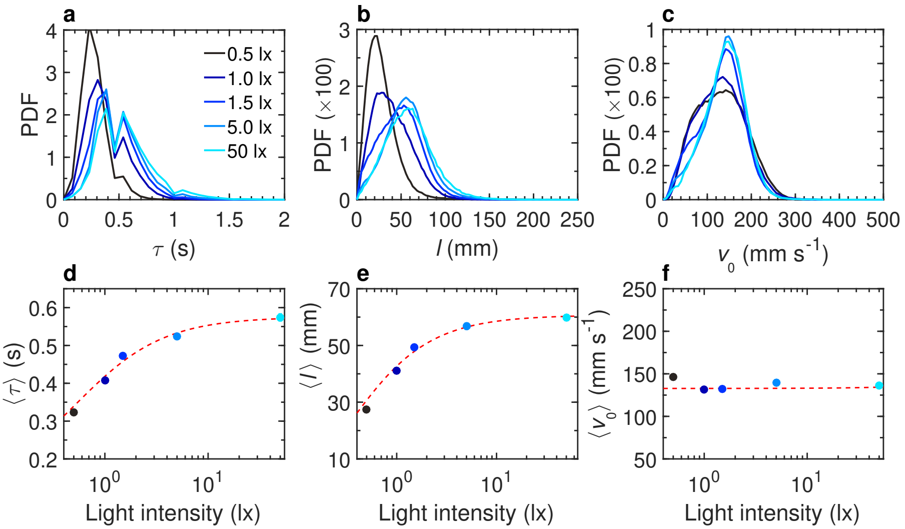


**Supplementary Fig. 11 Effects of light intensity on burst-and-coast swimming in groups of 25 fish a-c** Probability density function (PDF) of kick duration $\tau$, kick length $l$, and peak speed $v_0$ respectively, at different light intensities: 0.5, 1, 1.5, 5 and 50 lx (from dark to light blue). **d-f** Average value of kick duration $\langle\tau\rangle$, kick length $\langle l\rangle$, and peak speed $\langle v_0\rangle$ respectively, at different light intensities. Solid circles are the average values on all experiments; error bars represent the standard deviation. Red dashed lines show the trend of the average value with the light intensity.

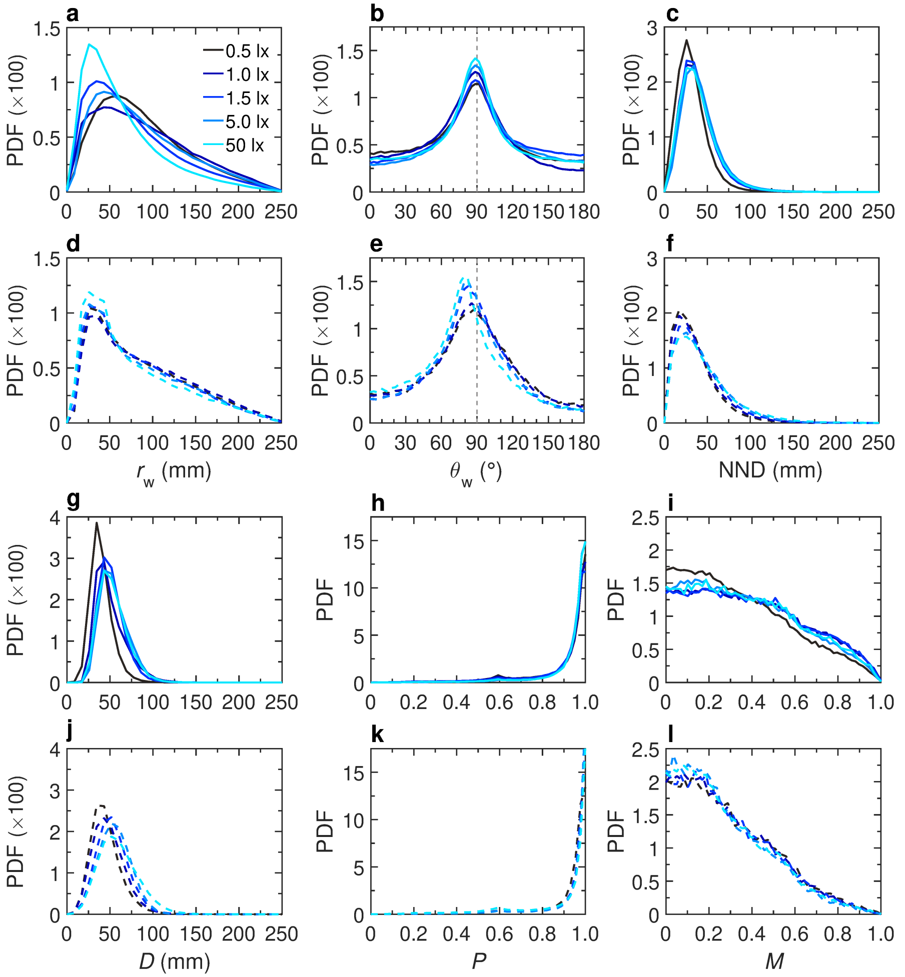


**Supplementary Fig. 12 Quantification of collective behavior in groups of 5 fish.** Probability density functions (PDF) of **a,d** the distance to the wall $r_w$, **b,e** the relative angle to the wall $\theta_w$, **c,f** the distance to the nearest neighbor NND, **g,j** dispersion $D$, **h,k** polarization $P$, and **i,l** milling $M$, for five different light intensities 0.5, 1, 1.5, 5, and 50 lx (from dark to light blue). Solid lines (**a-c, g-i**) correspond to experimental measures, dashed lines (**d-f, j-l**) to numerical simulations of the model.

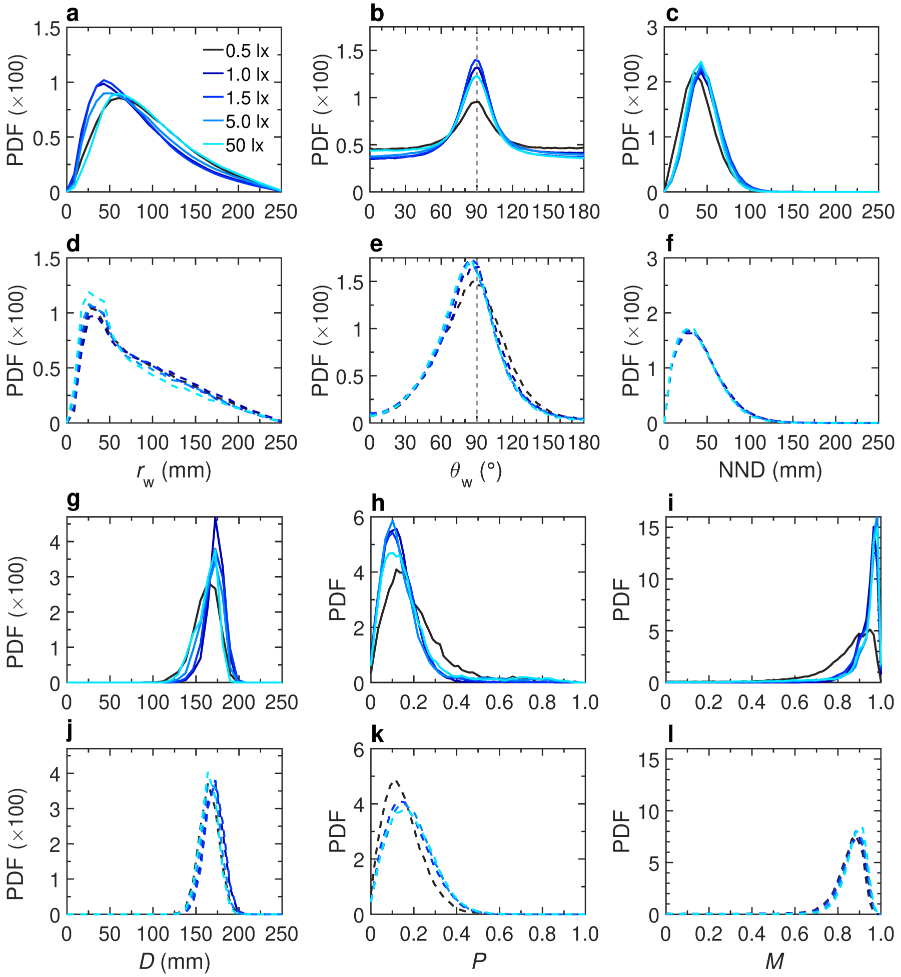


**Supplementary Fig. 13 Quantification of collective behavior in groups of 25 fish.** Probability density functions (PDF) of **a,d** the distance to the wall $r_w$, **b,e** the relative angle to the wall $\theta_w$, **c,f** the distance to the nearest neighbor NND, **g,j** dispersion $D$, **h,k** polarization $P$, and **i,l** milling $M$, for five different light intensities 0.5, 1, 1.5, 5, and 50 lx (from dark to light blue). Solid lines (**a-c, g-i**) correspond to experimental measures, dashed lines (**d-f, j-l**) to numerical simulations of the model.

### Supplementary Tables

**Supplementary Table 1** List of experiments with one fish.

| Light intensity | Date | Proportion of active swimming (%) | Duration (min) | #kicks |
|---|---|---|---|---|
| 0.5 lx | 2022-03-18 | 79.48 | 61 | 7960 |
| | 2022-03-28 | 79.09 | 61 | 8296 |
| | 2022-03-28 | 98.19 | 63 | 12064 |
| | 2022-03-30 | 95.16 | 59 | 11301 |
| | 2022-04-01 | 81.37 | 60 | 8615 |
| | 2022-04-04 | 99.83 | 60 | 12080 |
| | 2022-05-18 | 99.89 | 59 | 11391 |
| | 2022-05-19 | 100 | 59 | 11186 |
| | 2022-05-19 | 99.98 | 47 | 7688 |
| | 2022-05-20 | 100 | 62 | 11459 |
| 1 lx | 2022-03-18 | 96.82 | 66 | 12563 |
| | 2022-03-25 | 96.31 | 59 | 10909 |
| | 2022-03-30 | 99.87 | 57 | 10006 |
| | 2022-05-17 | 100 | 62 | 11438 |
| | 2022-05-17 | 100 | 59 | 7542 |
| | 2022-05-18 | 100 | 59 | 10040 |
| | 2022-05-19 | 100 | 60 | 10764 |
| | 2022-05-19 | 100 | 61 | 5691 |
| | 2022-05-19 | 100 | 59 | 7818 |
| | 2022-05-20 | 100 | 63 | 9547 |
| 1.5 lx | 2022-03-18 | 100 | 62 | 7496 |
| | 2022-03-25 | 100 | 59 | 8957 |
| | 2022-04-04 | 100 | 59 | 10335 |
| | 2022-04-04 | 99.89 | 59 | 11071 |
| | 2022-04-06 | 100 | 60 | 7507 |
| | 2022-05-18 | 100 | 59 | 8468 |
| | 2022-05-18 | 100 | 61 | 8611 |
| | 2022-05-19 | 100 | 60 | 6424 |
| | 2022-05-20 | 100 | 61 | 7233 |
| | 2022-05-20 | 100 | 58 | 6416 |
| 5 lx | 2022-03-09 | 100 | 61 | 7068 |
| | 2022-03-24 | 99.73 | 58 | 6388 |
| | 2022-04-06 | 100 | 61 | 5915 |
| | 2022-04-06 | 100 | 59 | 6398 |
| | 2022-05-17 | 100 | 57 | 8406 |
| | 2022-05-17 | 100 | 62 | 6743 |
| | 2022-05-18 | 100 | 60 | 4861 |
| | 2022-05-18 | 100 | 53 | 7553 |
| | 2022-05-19 | 100 | 60 | 7682 |
| | 2022-05-19 | 100 | 57 | 6084 |
| 50 lx | 2022-03-09 | 100 | 59 | 7156 |
| | 2022-03-24 | 100 | 58 | 6576 |
| | 2022-03-28 | 100 | 58 | 6972 |
| | 2022-03-28 | 100 | 59 | 5843 |
| | 2022-04-01 | 100 | 59 | 7752 |
| | 2022-05-17 | 100 | 61 | 6713 |
| | 2022-05-18 | 100 | 62 | 5913 |
| | 2022-05-18 | 100 | 58 | 8065 |
| | 2022-05-19 | 100 | 60 | 5934 |
| | 2022-05-19 | 100 | 65 | 7908 |

**Supplementary Table 2** List of experiments with two fish.

| Light intensity | Date | Proportion of active swimming (%) | Duration (min) | Number of kicks |
|---|---|---|---|---|
| 0.5 lx | 2022-03-07 | 96.84 | 56 | 19523 |
| | 2022-03-25 | 99.83 | 56 | 21319 |
| | 2022-03-29 | 96.44 | 58 | 18711 |
| | 2022-03-29 | 92.18 | 58 | 15798 |
| | 2022-04-01 | 99.32 | 57 | 21295 |
| | 2022-04-06 | 99.31 | 66 | 24590 |
| 1 lx | 2022-03-07 | 64.99 | 54 | 9826 |
| | 2022-03-25 | 99.61 | 59 | 19210 |
| | 2022-03-25 | 99.71 | 57 | 16479 |
| | 2022-03-29 | 97.90 | 59 | 20266 |
| | 2022-04-04 | 98.83 | 60 | 17000 |
| | 2022-05-20 | 99.61 | 59 | 19653 |
| 1.5 lx | 2022-03-11 | 97.43 | 56 | 14697 |
| | 2022-03-24 | 53.70 | 57 | 7680 |
| | 2022-03-25 | 96.18 | 58 | 14731 |
| | 2022-03-29 | 95.20 | 58 | 13974 |
| | 2022-04-04 | 98.34 | 61 | 14543 |
| | 2022-05-20 | 99.08 | 59 | 14797 |
| 5 lx | 2022-03-11 | 93.34 | 58 | 12219 |
| | 2022-03-24 | 69.91 | 59 | 8153 |
| | 2022-03-28 | 69.70 | 57 | 7133 |
| | 2022-03-29 | 95.66 | 59 | 10789 |
| | 2022-04-06 | 97.60 | 59 | 13426 |
| | 2022-05-20 | 40.13 | 61 | 3825 |
| 50 lx | 2022-03-07 | 65.00 | 54 | 9826 |
| | 2022-03-28 | 70.05 | 56 | 6387 |
| | 2022-03-29 | 54.29 | 59 | 4694 |
| | 2022-04-01 | 84.28 | 62 | 10107 |
| | 2022-04-06 | 67.46 | 58 | 5366 |
| | 2022-05-20 | 54.39 | 60 | 4968 |

**Supplementary Table 3** List of experiments with 5 fish.

| Light intensity | Date | Proportion of active swimming (%) | Duration (min) | Number of kicks |
|---|---|---|---|---|
| 0.5 lx | 2022-03-18 | 90.49 | 57 | 37817 |
| | 2022-03-18 | 88.42 | 57 | 28999 |
| | 2022-03-30 | 75.93 | 61 | 14714 |
| | 2022-04-01 | 66.21 | 55 | 12121 |
| | 2022-04-01 | 75.64 | 61 | 10760 |
| | 2022-04-05 | 80.20 | 55 | 18534 |
| 1 lx | 2022-03-08 | 97.94 | 60 | 42062 |
| | 2022-03-18 | 87.68 | 56 | 19928 |
| | 2022-03-18 | 89.32 | 57 | 25516 |
| | 2022-03-30 | 83.66 | 57 | 23991 |
| | 2022-04-04 | 85.14 | 59 | 16770 |
| 1.5 lx | 2022-03-16 | 85.64 | 56 | 17224 |
| | 2022-03-17 | 69.86 | 17 | 2193 |
| | 2022-03-28 | 80.94 | 63 | 13034 |
| | 2022-03-30 | 82.12 | 59 | 14797 |
| | 2022-04-04 | 85.67 | 58 | 14947 |
| 5 lx | 2022-03-16 | 88.76 | 56 | 16195 |
| | 2022-03-17 | 89.08 | 56 | 17241 |
| | 2022-03-28 | 87.03 | 60 | 15533 |
| | 2022-03-29 | 85.94 | 66 | 16616 |
| | 2022-04-05 | 79.12 | 60 | 11275 |
| 50 lx | 2022-03-17 | 84.27 | 57 | 11910 |
| | 2022-03-29 | 80.02 | 59 | 12131 |
| | 2022-03-30 | 84.62 | 56 | 11790 |
| | 2022-04-01 | 85.48 | 60 | 12728 |
| | 2022-04-01 | 73.50 | 54 | 8329 |

**Supplementary Table 4** List of experiments with 25 fish.

| Light intensity | Date | Proportion of active swimming (%) | Duration (min) | Number of kicks |
|---|---|---|---|---|
| 0.5 lx | 2022-03-09 | 68.77 | 58 | 90411 |
| | 2022-03-28 | 48.30 | 59 | 6645 |
| | 2022-03-31 | 46.02 | 60 | 5293 |
| | 2022-04-01 | 61.79 | 59 | 42853 |
| | 2022-04-04 | 39.21 | 58 | 3159 |
| | 2022-04-04 | 60.91 | 60 | 21468 |
| 1 lx | 2022-03-09 | 76.27 | 58 | 103074 |
| | 2022-03-25 | 73.88 | 59 | 76665 |
| | 2022-03-28 | 66.33 | 60 | 21493 |
| | 2022-03-31 | 43.12 | 59 | 15920 |
| | 2022-04-05 | 63.13 | 59 | 46706 |
| 1.5 lx | 2022-03-16 | 41.05 | 61 | 29354 |
| | 2022-03-25 | 58.63 | 60 | 34778 |
| | 2022-03-29 | 71.56 | 59 | 68492 |
| | 2022-03-31 | 68.86 | 59 | 68264 |
| | 2022-04-05 | 71.92 | 60 | 61727 |
| 5 lx | 2022-03-16 | 36.46 | 56 | 19369 |
| | 2022-03-18 | 14.18 | 53 | 3216 |
| | 2022-03-29 | 79.05 | 60 | 71719 |
| | 2022-03-31 | 63.51 | 56 | 33464 |
| | 2022-04-05 | 68.86 | 57 | 45622 |
| 50 lx | 2022-03-18 | 11.94 | 54 | 1826 |
| | 2022-03-31 | 50.57 | 58 | 14032 |
| | 2022-04-01 | 72.31 | 60 | 68599 |
| | 2022-04-04 | 17.89 | 60 | 2050 |
| | 2022-04-06 | 62.29 | 54 | 32073 |

**Supplementary Table 5** Parameters used in the simulations for $N = 1$.

| Parameters \ Light intensity(lx) | | 0.5 | 1 | 1.5 | 5 | 50 |
|---|---|---|---|---|---|---|
| $\gamma_R$ | Intensity of heading random fluctuations | 0.25 | 0.28 | 0.33 | 0.35 | 0.37 |
| $\gamma_w$ | Intensity of wall repulsion | 0.29 | 0.31 | 0.35 | 0.36 | 0.37 |
| $\tau_0$ | Relaxation time (s) | 0.34 | 0.66 | 0.76 | 0.83 | 0.87 |
| $\alpha$ | Fluctuations reduction factor when close to wall | 0.35 | 0.47 | 0.50 | 0.63 | 0.67 |
| $l_w$ | Range of wall repulsion (mm) | 31 | 45 | 50 | 52 | 53 |
| | Normalization constant of $O_w$ | 1.96 | 1.96 | 1.96 | 1.96 | 1.96 |
| $l_c$ | Comfort length (mm) | 30 | 30 | 30 | 30 | 30 |

**Supplementary Table 6** Parameters used in the simulations for $N = 2$.

| Parameters | | **Light intensity(lx)** **0.5** | **1** | **1.5** | **5** | **50** |
|---|---|---|---|---|---|---|
| $\gamma_{\mathrm{R}}$ | Intensity of heading random fluctuations | 0.25 | 0.28 | 0.33 | 0.35 | 0.37 |
| $\gamma_{\mathrm{w}}$ | Intensity of wall repulsion | 0.25 | 0.24 | 0.22 | 0.22 | 0.21 |
| $\gamma_{\mathrm{Att}}$ | Intensity of attraction/repulsion | 0.15 | 0.20 | 0.24 | 0.240 | 0.25 |
| $\gamma_{\mathrm{Ali}}$ | Intensity of alignment | 0.04 | 0.038 | 0.035 | 0.031 | 0.03 |
| $\tau_0$ | Relaxation time (s) | 0.34 | 0.66 | 0.76 | 0.83 | 0.87 |
| $\alpha$ | Fluctuations reduction factor when close to wall | 0.35 | 0.47 | 0.50 | 0.63 | 0.67 |
| $l_{\mathrm{w}}$ | Range of wall repulsion (mm) | 31 | 45 | 50 | 52 | 53 |
| $l_{\mathrm{Att}}$ | Range of attraction between individuals (mm) | 100 | 120 | 135 | 145 | 170 |
| $l_{\mathrm{Ali}}$ | Range of alignment between individuals (mm) | 92 | 105 | 115 | 125 | 145 |
| $d_{\mathrm{Att}}$ | Balance distance of attraction/repulsion (mm) | 15 | 15 | 15 | 15 | 15 |
| | Normalization constant of $O_{\mathrm{Att}}$ | 1.43 | 1.43 | 1.43 | 1.43 | 1.43 |
| | Normalization constant of $E_{\mathrm{Att}}$ | 0.887 | 0.887 | 0.887 | 0.887 | 0.887 |
| $d_{\mathrm{Ali}}$ | Balance distance of attraction/repulsion (mm) | 15 | 15 | 15 | 15 | 15 |
| | Normalization constant of $O_{\mathrm{Ali}}$ | 1.43 | 1.43 | 1.43 | 1.43 | 1.43 |
| | Normalization constant of $E_{\mathrm{Ali}}$ | 0.9 | 0.9 | 0.9 | 0.9 | 0.9 |
| $l_c$ | Comfort length (mm) | 35 | 35 | 35 | 35 | 35 |

**Supplementary Table 7** Parameters used in the simulations for $N = 5$.

| Parameters | Light intensity(lx) | **0.5** | **1** | **1.5** | **5** | **50** |
|---|---|---|---|---|---|---|
| $\gamma_{\mathrm{R}}$ | Intensity of heading random fluctuations | 0.15 | 0.19 | 0.21 | 0.22 | 0.25 |
| $\gamma_{\mathrm{w}}$ | Intensity of wall repulsion | 0.40 | 0.41 | 0.45 | 0.46 | 0.47 |
| $\gamma_{\mathrm{Att}}$ | Intensity of attraction/repulsion | 0.053 | 0.061 | 0.067 | 0.075 | 0.081 |
| $\gamma_{\mathrm{Ali}}$ | Intensity of alignment | 0.058 | 0.061 | 0.063 | 0.067 | 0.069 |
| $\tau_0$ | Relaxation time (s) | 0.34 | 0.66 | 0.76 | 0.83 | 0.87 |
| $\alpha$ | Fluctuations reduction factor when close to wall | 0.35 | 0.47 | 0.50 | 0.63 | 0.67 |
| $l_{\mathrm{w}}$ | Range of wall repulsion (mm) | 31 | 45 | 50 | 52 | 53 |
| $l_{\mathrm{Att}}$ | Range of attraction between individuals (mm) | 100 | 120 | 135 | 145 | 170 |
| $l_{\mathrm{Ali}}$ | Range of alignment between individuals (mm) | 92 | 105 | 115 | 125 | 145 |
| $d_{\mathrm{Att}}$ | Balance distance of attraction/repulsion (mm) | 15 | 15 | 15 | 15 | 15 |
| | Normalization constant of $O_{\mathrm{Att}}$ | 1.43 | 1.43 | 1.43 | 1.43 | 1.43 |
| | Normalization constant of $E_{\mathrm{Att}}$ | 0.887 | 0.887 | 0.887 | 0.887 | 0.887 |
| $d_{\mathrm{Ali}}$ | Balance distance of attraction/repulsion (mm) | 15 | 15 | 15 | 15 | 15 |
| | Normalization constant of $O_{\mathrm{Ali}}$ | 1.43 | 1.43 | 1.43 | 1.43 | 1.43 |
| | Normalization constant of $E_{\mathrm{Ali}}$ | 0.9 | 0.9 | 0.9 | 0.9 | 0.9 |
| $l_c$ | Comfort length (mm) | 45 | 45 | 45 | 45 | 45 |

**Supplementary Table 8** Parameters used in the simulations for N=25.

| Parameters | Light intensity(lx) | **0.5** | **1** | **1.5** | **5** | **50** |
|---|---|---|---|---|---|---|
| $\gamma_{\mathrm{R}}$ | Intensity of heading random fluctuations | 0.02 | 0.08 | 0.1 | 0.08 | 0.02 |
| $\gamma_{\mathrm{w}}$ | Intensity of wall repulsion | 0.29 | 0.31 | 0.35 | 0.36 | 0.37 |
| $\gamma_{\mathrm{Att}}$ | Intensity of attraction/repulsion | 0.02 | 0.02 | 0.02 | 0.018 | 0.017 |
| $\gamma_{\mathrm{Ali}}$ | Intensity of alignment | 0.004 | 0.007 | 0.008 | 0.008 | 0.007 |
| $\tau_0$ | Relaxation time (s) | 0.34 | 0.66 | 0.76 | 0.83 | 0.87 |
| $\alpha$ | Fluctuations reduction factor when close to wall | 0.35 | 0.47 | 0.50 | 0.63 | 0.67 |
| $l_{\mathrm{w}}$ | Range of wall repulsion (mm) | 31 | 45 | 50 | 52 | 53 |
| $l_{\mathrm{Att}}$ | Range of attraction between individuals (mm) | 100 | 120 | 135 | 145 | 170 |
| $l_{\mathrm{Ali}}$ | Range of alignment between individuals (mm) | 92 | 105 | 115 | 125 | 145 |
| $d_{\mathrm{Att}}$ | Balance distance of attraction/repulsion (mm) | 15 | 15 | 15 | 15 | 15 |
| | Normalization constant of $O_{\mathrm{Att}}$ | 1.43 | 1.43 | 1.43 | 1.43 | 1.43 |
| | Normalization constant of $E_{\mathrm{Att}}$ | 0.887 | 0.887 | 0.887 | 0.887 | 0.887 |
| $d_{\mathrm{Ali}}$ | Balance distance of attraction/repulsion (mm) | 15 | 15 | 15 | 15 | 15 |
| | Normalization constant of $O_{\mathrm{Ali}}$ | 1.43 | 1.43 | 1.43 | 1.43 | 1.43 |
| | Normalization constant of $E_{\mathrm{Ali}}$ | 0.9 | 0.9 | 0.9 | 0.9 | 0.9 |
| $l_c$ | Comfort length (mm) | 30 | 30 | 30 | 30 | 30 |

### Supplementary Movies

**- Video S1: Effect of light intensity on individual swimming behavior in rummy-nose tetra (***Hemigrammus rhodostomus).*

Video excerpts of experiments with a single fish swimming alone in a circular tank of radius 250 mm under five different light intensities (0.5, 1, 1.5, 5, and 50 lx).

**- Video S2: Effect of light intensity on social interactions between two fish in rummy-nose tetra (***Hemigrammus rhodostomus).*

Video excerpts of experiments with 2 fish swimming in a circular tank of radius 250 mm under five different light intensities (0.5, 1, 1.5, 5, and 50 lx).

**- Video S3: Effect of light intensity on collective behavior in groups of 5 fish in rummy-nose tetra (***Hemigrammus rhodostomus).*

Video excerpts of experiments with a group of 5 fish swimming in a circular tank of radius 250 mm under five different light intensities (0.5, 1, 1.5, 5, and 50 lx).

**- Video S4: Effect of light intensity on collective behavior in groups of 25 fish in rummy-nose tetra (***Hemigrammus rhodostomus).*

Video excerpts of experiments with a group of 25 fish swimming in a circular tank of radius 250 mm under five different light intensities (0.5, 1, 1.5, 5, and 50 lx).

**- Video S5: Numerical simulations of the model with a single fish under different light intensities.**

Representative example of a simulation of a single fish swimming in a circular tank of radius 250 mm under five different light intensities (0.5, 1, 1.5, 5, and 50 lx). The size of the simulated fish does not correspond to the actual dimensions of the real fish and is used for ease of visualization.

**- Video S6: Numerical simulations of the model with 2 fish under different light intensities.**

Representative example of a simulation of 2 fish swimming in a circular tank of radius 250 mm under five different light intensities (0.5, 1, 1.5, 5, and 50 lx). The size of the simulated fish does not correspond to the actual dimensions of the real fish and is used for ease of visualization.

**- Video S7: Numerical simulations of the model with a group of 5 fish under different light intensities.**

Representative example of a simulation of a group of 5 fish swimming in a circular tank of radius 250 mm under five different light intensities (0.5, 1, 1.5,

5, and 50 lx). Each fish interacts with its two most influential neighbors. The size of the simulated fish does not correspond to the actual dimensions of the real fish and is used for ease of visualization.

**- Video S8: Numerical simulations of the model with a group of 25 fish under different light intensities.**

Representative example of a simulation of a group of 25 fish swimming in a circular tank of radius 250 mm under five different light intensities (0.5, 1, 1.5, 5, and 50 lx). Each fish interacts with its two most influential neighbors. The size of the simulated fish does not correspond to the actual dimensions of the real fish and is used for ease of visualization.

### Supplementary Data

***Supplementary Data 1:***

All data needed to evaluate and replicate the conclusions of the article are present in the article, the Supplementary Materials, or available at the following online repository: https://doi.org/10.6084/m9.figshare.22640053.v1.
